# Geographically Weighted Linear Combination Test for Gene Set Analysis of a Continuous Spatial Phenotype as applied to Intratumor Heterogeneity

**DOI:** 10.1101/2022.10.09.511477

**Authors:** Payam Amini, Morteza Hajihosseini, Saumyadipta Pyne, Irina Dinu

**Affiliations:** Department of Biostatistics, School of Public Health, Iran University of Medical Sciences, Tehran, Iran; School of Public Health, University of Alberta, Edmonton, AB, Canada; Health Analytics Network, Pittsburgh, PA, USA; Department of Statistics and Applied Probability, University of California, Santa Barbara, CA, USA

**Keywords:** Intratumor heterogeneity, Gene-set analysis, Geographically Weighted Regression, Linear Combination Test, Spatial single cell analysis, Cancer-associated fibroblasts

## Abstract

**Background:** The impact of gene-sets on phenotype is not necessarily uniform across different locations of a cancer tissue. This study introduces a computational platform, GWLCT, for combining gene set analysis with spatial data modeling to provide a new statistical test for association of phenotypes and molecular pathways in spatial single-cell RNA-seq data collected from an input tumor sample.

**Methods:** At each location, the most significant linear combination is found using a geographically weighted shrunken covariance matrix and kernel function. Whether a fixed or adaptive bandwidth is determined based on a cross validation procedure. Our proposed method is compared to the global version of linear combination test (LCT), bulk and random-forest based gene-set enrichment analyses using data created by the Visium Spatial Gene Expression technique on an invasive breast cancer tissue sample, as well as 144 different simulation scenarios.

**Results:** In an illustrative example, the new geographically weighted linear combination test, GWLCT, identifies the cancer hallmark gene-sets that are significantly associated at each location with the five spatially continuous phenotypic contexts in the tumors defined by different well-known markers of cancer-associated fibroblasts. Scan statistics revealed clustering in the number of significant gene-sets. A spatial heatmap of combined significance over all selected gene-sets is also produced. Extensive simulation studies demonstrate that our proposed approach outperforms other methods in the considered scenarios, especially when the spatial association increases.

**Conclusions:** Our proposed approach considers the spatial covariance of gene expression to detect the most significant gene-sets affecting a continuous phenotype. It reveals spatially detailed information in tissue space and can thus play a key role in understanding contextual heterogeneity of cancer cells.

## Introduction

Extensive studies over the past few decades have uncovered a variety of cell populations in tumors, thus leading to the active research area of intratumor heterogeneity (ITH) (1). In 2010, Hanahan and Weinberg noted that tumors exhibit an additional dimension of complexity through their “tumor microenvironment” that contributes to the acquisition of the so-called hallmark traits of cancer. ITH is attributed to genetic, epigenetic, and microenvironmental factors (1, 2) and associated with poor prognosis, therapeutic resistance and treatment failure leading to poor overall survival in cancer patients (3–7). Indeed, the persistence of some of the drug-tolerant intratumor cell populations could be attributed to their high phenotypic plasticity (8).

Interestingly, hierarchies of differentiation also exist among normal cells in healthy tissues, but the populations of tumor cells display far greater cell-to-cell variability and the resulting phenotypic instability (9, 10). Such ITH could be attributed to genetic causes ranging from aneuploidy to spontaneous cell fusions, say, between cancer and non-cancer cells, in addition to other factors such as complex contextual signals in the highly aberrant tumor microenvironments, or even global alterations in cancer cell epigenomes (11). ITH also involves immune cell infiltration, which is of obvious importance for immunotherapies. Tumor antigen diversity could be determined by the T cell clonality in the different regions of the same tumor (11). Studies have shown spatially complex interactions between tumor microenvironments and the patient’s immune system (12, 13).

While heterogeneous cell types are prevalent within the tumor microenvironment some of which may account for cancer development and progression, it also contains different non-malignant components, including the cancer-associated fibroblasts (CAFs) (14–16). Although the origin and activation mechanism of CAFs remains an area of active research (17–20), studies have attributed the processes of formation and derivation of CAFs to various precursor cells (17–20), which may be the source of the well-known heterogeneity among the CAFs (21–24). Indeed, certain tumors, such as in the breast, in which the prevalence of CAFs could be as high as 80%, they can play both anti- as well as pro-tumorigenic roles (25–27). Importantly, CAFs can facilitate drug resistance dynamically by altering the cell-matrix interactions that control the outer layer of cells’ sensitivity to apoptosis, producing proteins that control cell survival and proliferation, assisting with cell-cell communications, and activating epigenetic plasticity in neighboring cells (28, 29). CAF-targeted treatments can have dual effects depending on the target and the tissue under consideration (30–32). For instance, spatial proximity to CAFs has been shown to impact molecular features and therapeutic sensitivity of breast cancer cells influencing clinical outcomes (33).

Globally, there were approximately 2.3 million new cases of and 685,000 deaths due to breast cancer (BC) in 2020 (34). BC is also the leading cause of cancer-related deaths among women (35, 36) and the second leading cause of cancer deaths globally, worldwide (35). The tumorigenesis involves uncontrolled growth of cells in breast tissues which can be either benign or malignant (37). Several studies on breast cancer patients have revealed the different anti- and pro-tumorigenic roles of the CAFs involved (25–27). Findings from various studies have also noted the functional heterogeneity of CAFs thus resulting in both diverse phenotypes in breast cancer and their increasing role in influencing the therapeutic outcomes (38).

In recent years, higher resolution, tissue-specific gene expression analysis is made possible by using new platforms such as single-cell RNA sequencing (scRNA-seq), which has rapidly evolved as a powerful and popular tool (39, 40). Unlike previous transcriptomic studies that assayed a “bulk” sample, scRNA-seq data can provide a detailed characterization of each tumor. Indeed, the Human Tumor Atlas Network [https://humantumoratlas.org] is increasingly enriched with data on human cancers based on scRNA-seq assays. The high-resolution transcriptomic platform has led to several scRNA-seq studies of the composition of CAFs in different stages of cancer (41–49). For focused understanding of the heterogeneous expressions of genes, different sites of the same tumor were analyzed with multiregional RNA sequencing for different cancers (7, 50–52).

Despite the advancements and efficacy of scRNA-seq, the lack of spatial information in scRNA-seq analysis is a significant shortcoming for typical scRNA-seq methods to capture cellular heterogeneity. For a tumor sample, the presence of spatial contexts might play a major role which could be combined with scRNA-seq data with the explicit aim to capture microenvironmental heterogeneity. Spatial cell-to-cell communication in a given tissue image can be recovered from a spatial scRNA sequencing data via computational spatial re-mapping (53). Alternatively, integration of high-resolution gene expression data with spatial coordinates can resolve such experimental shortcomings (54). While imaging the transcriptome in situ with high accuracy has been a major challenge in single-cell biology, development of high-throughput platforms for sequential fluorescence in situ hybridization such as RNA seqFISH+ and algorithms such as CELESTA can identify cell populations and their spatial organization in intact tissues (7, 55). Towards this, many recent efforts have developed methods to analyze spatial information in single-cell studies (56–64).

Be they generated by RNA-seq or microarrays in the past, while high-throughput transcriptomic data are useful for identifying genes that are differentially expressed, they are also used to test for co-regulation of multiple genes, i.e., a gene-set, based on existing empirical knowledge of biological pathways and gene signatures, e.g., the well-known hallmarks of cancer. In this direction, several methods for gene-set analysis (GSA) were introduced by (65–70). Since the genes within such gene-sets share a common biological function, considering the correlations within each set is a key aspect of a useful GSA method. However, it was shown by (71) that the above GSA methods were affected by large type II errors.

An important limitation of many GSA methods is that they can only accommodate binary outcomes, such as disease versus control. Our method, Linear Combination Test (LCT) is a GSA method that was designed to address these limitations by taking into account correlations across genes and outcomes, and dealing with binary, univariate or multivariate continuous outcomes, measured either at a single point in time or at multiple time points, and therefore, allow us to analyze a wider range of studies involving complex study designs (72). Studies have shown that LCT can overcome difficulties such as small sample size, large gene-sets, and can accommodate correlations across gene-sets, time points, and multiple correlated continuous phenotypes (73). Thus, while a specific gene may not show consistent expression across individual cells, LCT is more likely than traditional approaches to detect the regulation of a functional process or biological pathway associated with the intercellular diversity of outcomes in a single cell level experiment.

Recently, we have extended LCT beyond any other “bulk” GSA method for application to single cell experiments (74). However, GSA is considerably more complicated in the presence of spatial information since the analyzed gene-sets need not have a uniform impact over the entire area of a spatially continuous phenotype. In fact, the significance of association between a selected gene-set and a particular phenotypic context at various microenvironmental neighborhoods could be different. Yet, variable as they may be, since spatial effects are generally continuous in nature, proximity may determine more correlated associations than those across distant locations within the same tumor space. Notably, this alludes to Tobler’s First Law of Geography, which states that “everything is related to everything else, but near things are more related than distant things.”

Traditional testing of such relationships involves global or “aspatial” regression, with the implicit assumption that the impact of the genes in a gene-set (covariates) on the phenotype (spatial outcome) is constant across the tumor space (study area). In the presence of ITH, such stationarity assumption is unlikely to be valid. Geographically weighted regression (GWR) is a well-known method (75) that avoids this problem by performing the regression within local windows and each observation is weighted according to its proximity to the center of the window. Adaptive kernel bandwidths allow for heterogeneity among densities of gene expression over the windows in different parts of the study region. Local regression coefficients and associated statistics are mapped to visualize how the explanatory power of a gene-set on the associated phenotypes changes spatially.

In the present study, we combined gene-set analysis of LCT with the local spatial modeling of GWR with the aim to develop geographically weighted LCT (GWLCT) as a statistical test. We demonstrated it on spatial scRNA-seq data from a real breast tumor sample and obtained key insights into its molecular heterogeneity across different spatially continuous phenotypic contexts defined by five well-known markers of CAFs. We note that GWLCT has several distinct advantages. While the popular GSA methods are aspatial and use only bulk gene expression data, GWLCT is developed for spatial single cell gene expression data. The geographical weighting scheme allows nearby neighborhoods to contribute more to each local model, and the regions with significant association of a selected gene-set and a corresponding phenotype are detected using scan statistics on the local test scores and illustrated as maps. A spatial heatmap of combined significance over all gene-sets is produced. We also present new 3D interactive tools for insightful visualization of the tumor space. In the next section, we describe the data and methods, followed by the results of real tumor data analysis and simulations of different association scenarios using GWLCT, and end with discussion including future work.

## Materials and Methods

### Data

Data for spatial transcriptomics were downloaded from the 10x Genomics website [https://www.10xgenomics.com/]. In brief, the data were created using the Visium Spatial Gene Expression technique on an invasive breast cancer tissue sample that is expressing the Estrogen Receptor (ER), Progesterone Receptor (PR), and Human Epidermal Growth Factor Receptor (HER) negative. Illumina NovaSeq 6000 was used to generate the RNA sequencing data, which had a sequencing depth of 72,436 mean reads per cell. The downloaded dataset was filtered for average gene expression values greater than 1, and the resulting data matrix had 1,981 rows (genes) and 4,325 columns (single cells). The zero counts were substituted as part of the RNAseq data preparation with a relatively small random jitter about zero that would have the least impact on the remaining gene expression values. Using the bestNormalize package in the R programming language, we used a 10-fold cross-validation based data transformation strategy to normalize each gene’s expression across samples (76).

### Gene Sets

We downloaded from the Molecular Signatures Database (MSigDB) candidate gene-sets that represent commonly known “hallmarks” of cancer (77). To ensure their relevance as well as non-redundancy, we selected 8 of those hallmark gene-sets that have at least 25% overlap with the expressed genes (see above text on preprocessing) but mutual gene-set overlap of less than 10%. The selected hallmark gene-sets are: Epithelial Mesenchymal Transition (EMT, size= 81) (78), Angiogenesis (size=12) (79), DNA Repair (DNA_Rep, size=42) (80), Pi3k AKT MTOR Signaling (Pi3k, size=28) (81), Fatty Acid Metabolism (FAM, size=41) (82), P53 Pathway (P53, size=50) (83), Estrogen Response Early (ERE, size=63) (84), and Estrogen Response Late (ERL, size=62) (85).

### CAF Markers

A selected set of five CAF phenotypes, which were represented by the expression of the corresponding marker genes (the respective phenotypes are noted in parentheses): *CXCL12* (CAF-S1), *FBLN1* (mCAFs), *C3* (inflammatory CAFs), *S100A4* (sCAFs), and *COL11A1,* which is a fibroblast-specific “remarkable biomarker” that codes for collagen 11-α1 and shows expression gain in CAFs (86). For details on the CAF markers, see reviews, e.g., (87, 88).

### Statistical Analysis

Consider gene expression data of ‘*g*’ gene variables (*X*_1_, *X*_2_, *X*_3_, …, *X_g_*), ‘*L*’ cells (points) and ‘*K*’ sets of genes in a coordination format such as Cartesian. The LCT approach assumes a null hypothesis in which there is no association between a linear combination of *X*_1_, *X*_2_, *X*_3_, …, *X_g_* with the phenotype (73). For a local point (*u_l_, v_l_*), we can define a univariate regression as:

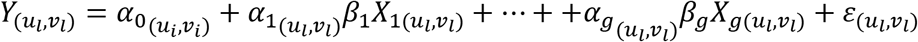

where

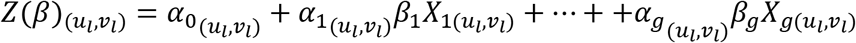

is the linear combination of *X*_1_, *X*_2_, *X*_3_, …, *X_g_* and 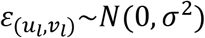. For each location in the dataset, we can find the most significant linear combination as follows:

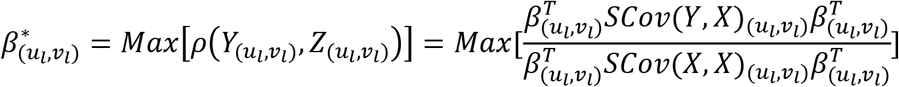

where *SCov* represents the weighted shrunken covariance matrix for each calibration location. The weights are generated using a bisquare kernel function, based on the Euclidean distance between two points *l* and *l’*

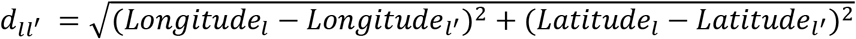

and bandwidth *h_l_*, which determines the radius around the point *l.* Here, the optimal bandwidth is calculated using cross validation (CV) based on the sum of squared errors at each cell point and set of genes:

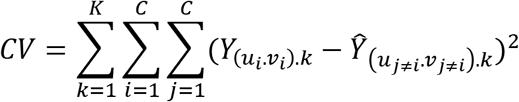

The bandwidth with the least measure of CV is used for localization. Weighting functions of bisquare and tricube type kernels are used to take the weighted location at *l* against another location *l’* into account. The bisquare kernel weighting function is defined as:

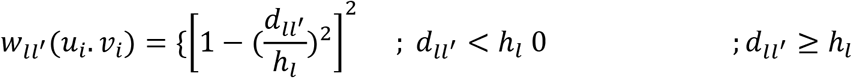

and the tricube kernel weighting function as:

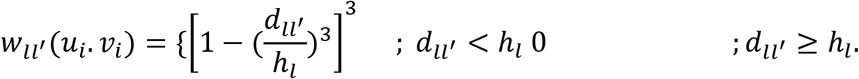

For the weighting functions, the bandwidth can be determined either beforehand (fixed distance) or as the distance between the point *l* and its nearest neighbor (adaptive), which is predetermined as well.

The shrunken covariance matrix of the gene expression data in the *l^th^* cell and around the estimated bandwidth *(h)* can be written as:

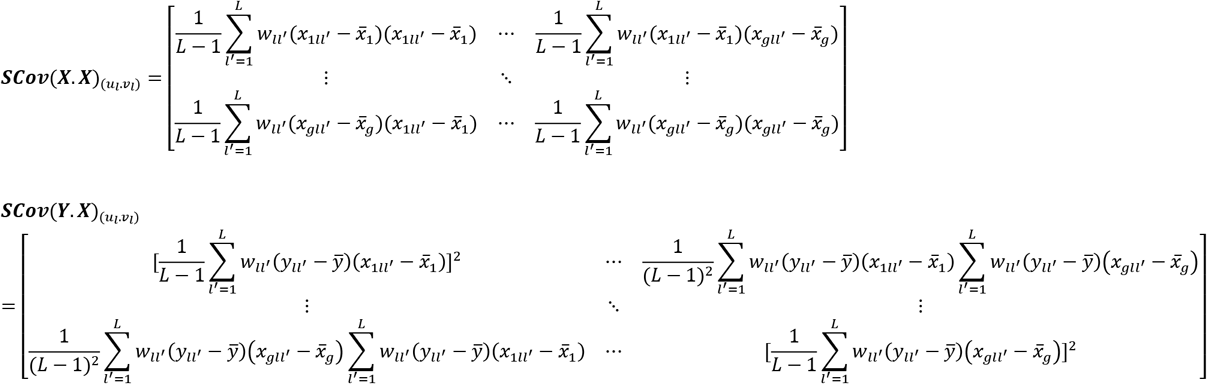

Using the weighted shrunken covariance matrices, the most significant linear combination at location (u_z_, *V_l_*) can be determined as the maximum Eigenvector of *SCov*(*Y,X*)_(*u_l_, v_l_*)_ *SCov*(*X,X*)_(*u_l_, v_l_*)_^-1^

### Combined significance mapping

We used the Fisher’s sum of log of independent p-values method to calculate the combined significance (CS) of association of the *K* gene-sets (e.g., 8 hallmarks in the present example) with a fixed phenotype at each location. The sum 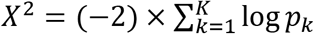 follows a chi-square distribution with 2*K* degrees of freedom, which yields a combined p-value *CP* corresponding to *X^2^* at a given location. Thus, the combined significance is computed as *CS* = (−1) × log*CP* and plotted as a spatial heatmap.

### Spatial cluster detection

Based on the count of significant gene-sets as determined by GWLCT at a given location, spatial clusters are detected and mapped for the user. For this purpose, assuming a Poisson distribution of such counts over a grid of points placed on the tumor space, scan statistics are computed with Openshaw’s Geographical Analysis Machine (GAM) (89) function as implemented in the R package DCluster.

### Comparative analysis

In addition to GWLCT, the global LCT, GSEA, and a Random Forest based GSEA (RF-GSEA) (90) were also performed to identify the global gene-sets that can accurately predict samples. In the domain of regression, the random-forest based technique is used when the outcome of concern is a continuous phenotype. The GSEA ignores the continuous phenotype and checks if the gene-sets show statistical difference between biological states. The RF-GSEA combines bootstrap and classification tree to find the proportion of explained variance of a continuous phenotype for a specific gene-set. The small sample size issue has been previously considered in these methods so that variable selection is conducted only from a small random subset of the variables. Moreover, the RF-GSEA is able to accommodate a continuous phenotype when the associations between genesets and phenotypes are nonlinear and contains complicated high-order interaction effects (90).

### Simulation study

Several scenarios were designed based on the potential impact of simulation components on the statistical power of the methods. Similar to our previous simulations studies for LCT and its extensions (73), we selected a random subset of genes from the expression data matrix, of a pre-defined size, i.e. the total number of genes used in each simulation experiment. Then we used spatial and spatio-temporal geostatistical modeling, prediction and simulation function in R software called ‘gstat’. This function creates an R object with the necessary fields for univariate or multivariate geostatistical prediction, its conditional or unconditional Gaussian, or indicator simulation equivalents (91).

Values were set for the variogram model components as 10 for the partial sill, 3 for the range parameter, 10 for the nugget, and 30 for the number of nearest observations that is used for kriging simulation. Moreover, a Gaussian model was assumed for the distribution of the gene expressions and phenotypes. In the next step, the GWLCT components were defined. For the gene-set matrix, a binomial distribution was used to generate a membership indicator matrix in which the proportions of genes belonging to the gene-sets were characterized using the probability parameter (Low = 0.3, and High = 0.9). Three different values for the spatial covariance were considered to imply the spatial association among genes’ expressions. The higher the spatial covariance, the lower is the spatial association. Thus, a variance of 50 was considered as high which gives a corresponding low spatial association, a variance of 5 a moderate spatial association, and a variance of 0.1 a high spatial association.

As well, the spatial association between the continuous phenotype and the gene expression data was taken into account by a spatially and normally distributed phenotype generated from the gene expression data with the same parameters as for the spatial association among genes’ expressions. High and low levels for the radius/ bandwidth around each location for the local analyses were 20 and 6 respectively. Number of coordinate points was 10 by 10 for low, and 100 by 100 for high. Finally, the two levels of low and high were defined for the total number of genes as 100 and 1000 respectively.

The above simulation components resulted in 144 scenarios in which an adaptive bandwidth kernel with the bisquare weighting function and were used. The number of permutations was fixed and considered as 500 and the threshold of significance was assumed as 0.05. The methods were compared based on their statistical power. An average of statistical powers across the locations was computed and used as the performance measure for GWLCT.

R programming software version 4.1.1 is used for data analysis and packages such as corpcor (92), qvcalc (93), stringr (94), and plotly (95).

## Results

The three aspatial methods, LCT, RF-GSEA, and GSEA, were used to identify the “hallmark” cancer gene-sets that are significantly associated with five spatially continuous CAF phenotypes represented by their known markers C3, COL11A1, CXCL12, FBLN1, and S100A4 in the single cell breast cancer spatial transcriptomic data. The three aspatial methods, the phenotype, the size of each gene-set, p-value of the test, and the corresponding q-value are shown in Table 1. We note that such global p-values cannot be obtained from our proposed GWLCT, as this method is spatial in nature and assesses significance at every coordinate point, rather than an overall significance measure. Using GSEA, the only significant gene-set was P53 (p-value=0.009, q-value=0.010). The results of LCT and RF-GSEA revealed that all the gene-sets are strongly associated with the five continuous phenotypes with p-value and q-value less than 0.001. This was expected since the candidate gene-sets represent commonly known “hallmarks” of cancer (77).

**Table 1.** An evaluation of cancer hallmark gene-sets associated with five continuous phenotypes C3, COL11A1, CXCL12, FBLN1, and S100A4 using the aspatial methods including GSEA, LCT, and RF-GSEA on a single cell breast cancer study.

| Method | Phenotype | Gene-set name | Gene-set size | P-value | Q-value |
| --- | --- | --- | --- | --- | --- |
| GSEA (No phenotype specified) | Not Applicable | EMT | 81 | 0.504 | 0.648 |
|  |  | ANGIOGENESIS | 12 | 0.227 | 0.486 |
|  |  | DNA_Rep | 41 | 0.425 | 0.763 |
|  |  | PI3K | 28 | 0.059 | 0.155 |
|  |  | FAM | 40 | 0.623 | 0.648 |
|  |  | P53 | 50 | 0.009 | 0.010 |
|  |  | ERE | 64 | 0.178 | 0.486 |
|  |  | ERL | 62 | 0.297 | 0.486 |
| Global LCT | C3, COL11A1, CXCL12, FBLN1, S100A4 | EMT | 81 | <0.001 | <0.001 |
|  |  | ANGIOGENESIS | 12 | <0.001 | <0.001 |
|  |  | DNA_Rep | 41 | <0.001 | <0.001 |
|  |  | PI3K | 28 | <0.001 | <0.001 |
|  |  | FAM | 40 | <0.001 | <0.001 |
|  |  | P53 | 50 | <0.001 | <0.001 |
|  |  | ERE | 64 | <0.001 | <0.001 |
|  |  | ERL | 62 | <0.001 | <0.001 |
| RF-GSEA | C3, COL11A1, CXCL12, FBLN1, S100A4 | EMT | 81 | <0.001 | <0.001 |
|  |  | ANGIOGENESIS | 12 | <0.001 | <0.001 |
|  |  | DNA_Rep | 41 | <0.001 | <0.001 |
|  |  | PI3K | 28 | <0.001 | <0.001 |
|  |  | FAM | 40 | <0.001 | <0.001 |
|  |  | P53 | 50 | <0.001 | <0.001 |
|  |  | ERE | 64 | <0.001 | <0.001 |
|  |  | ERL | 62 | <0.001 | <0.001 |

Scan statistics were computed to detect the potential clusters of spatial regulation based on the number of significant gene-sets at the locations where such counts exceeded what may be expected from an underlying Poisson distribution over the tumor space (Figure 1). For the local GWLCT, three different CAF categories were identified as Low (CAF gene expression less than 0.5), Moderate (CAF gene expression between 0.5 and 1), and High (CAF gene expression more than 1). The 3D plots are presented here as well as at the following links on GitHub: https://mortezahaji.github.io/GWLCT-Project/. By clicking on any point in the plot, one can find the coordinate of the cell, corresponding number and name of significant gene-sets at three different levels of Low, Moderate, and High CAF gene levels.

**Figure 1.**
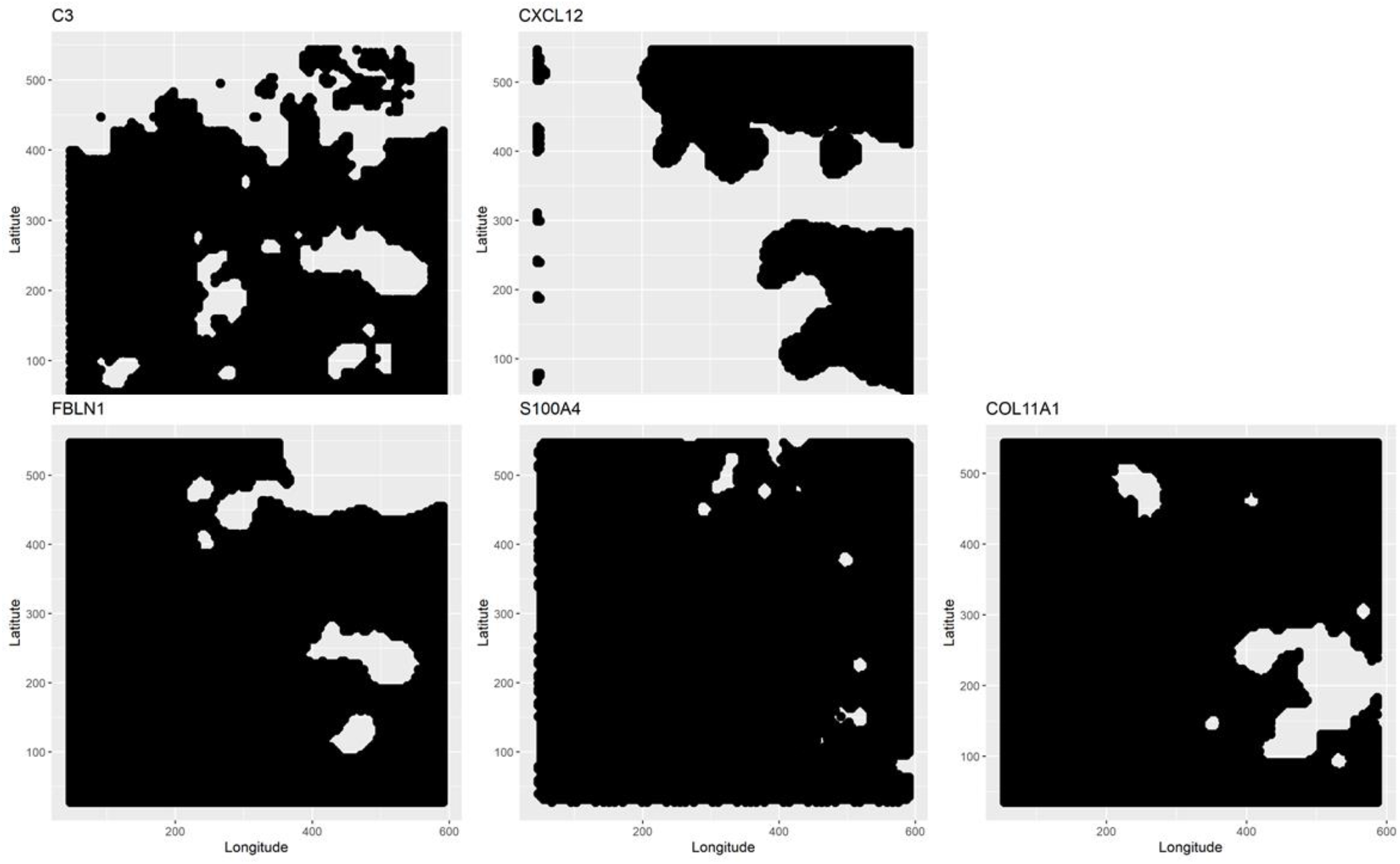
Scan statistics to detect and demarcate the potential clusters based on the number of significant gene-sets following a Poisson distribution

**Figure 2.**
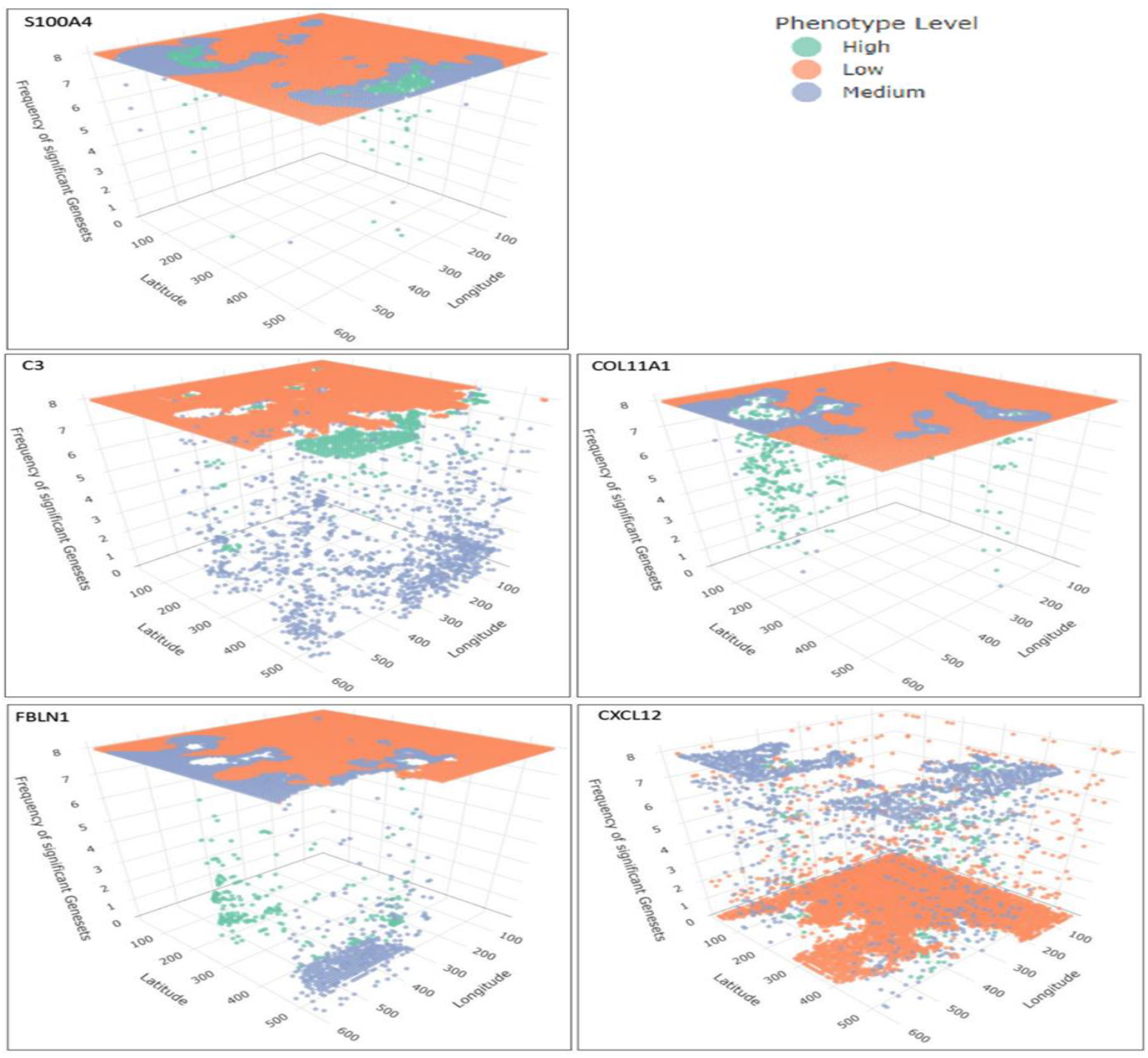
demonstrates a snapshot of the 3D plots for the 5 phenotypes at three CAF levels. A snapshot of the 3D plots for the 5 phenotypes at Low, Medium and High CAF levels. The interactive plots can be accessed at: https://mortezahaji.github.io/GWLCT-Project/

A snapshot of the 3D plot for the phenotype COL11A1 at high CAF level is also demonstrated in Figure 3.

**Figure 3.**
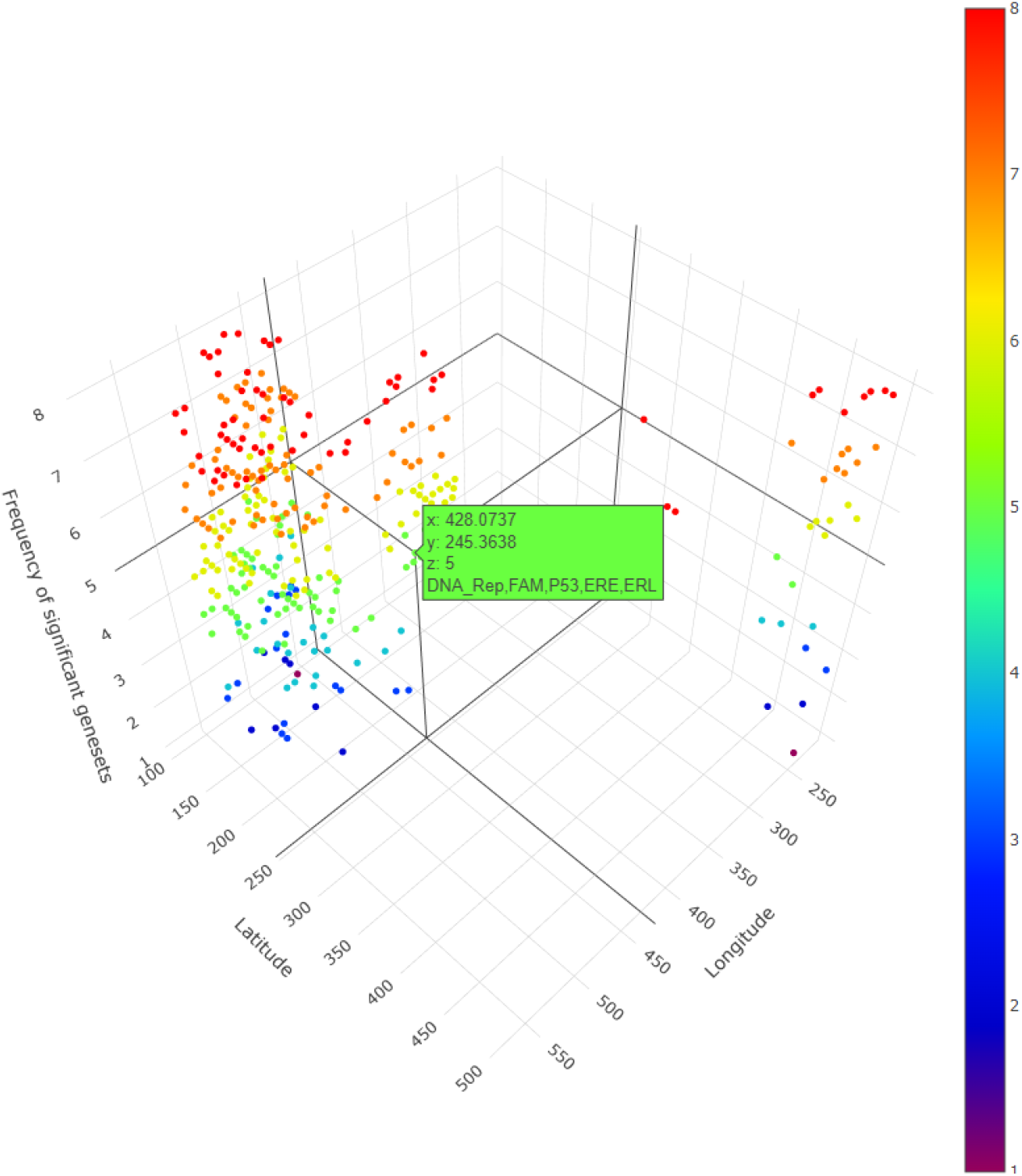
A snapshot of the 3D plot for COL11A1 at high CAF level. The number of significant gene-sets are shown in 8 colors (based on 8 gene-sets). The interactive plots can be accessed at: https://mortezahaji.github.io/GWLCT-Project/

One is able to detect the frequency as well as the names of significant gene-sets at each location (based on the 8 gene-sets) by clicking on each dot, rotating and zooming into the 3D interactive plot available at: https://mortezahaji.github.io/GWLCT-Project/. The 3D plot in Figure 3 is divided into eight 2-dimension plots in Figure 4 so that one can evaluate the distribution of one to eight significant gene-sets across the regions with COL11A1 expressed at high CAF level.

**Figure 4.**
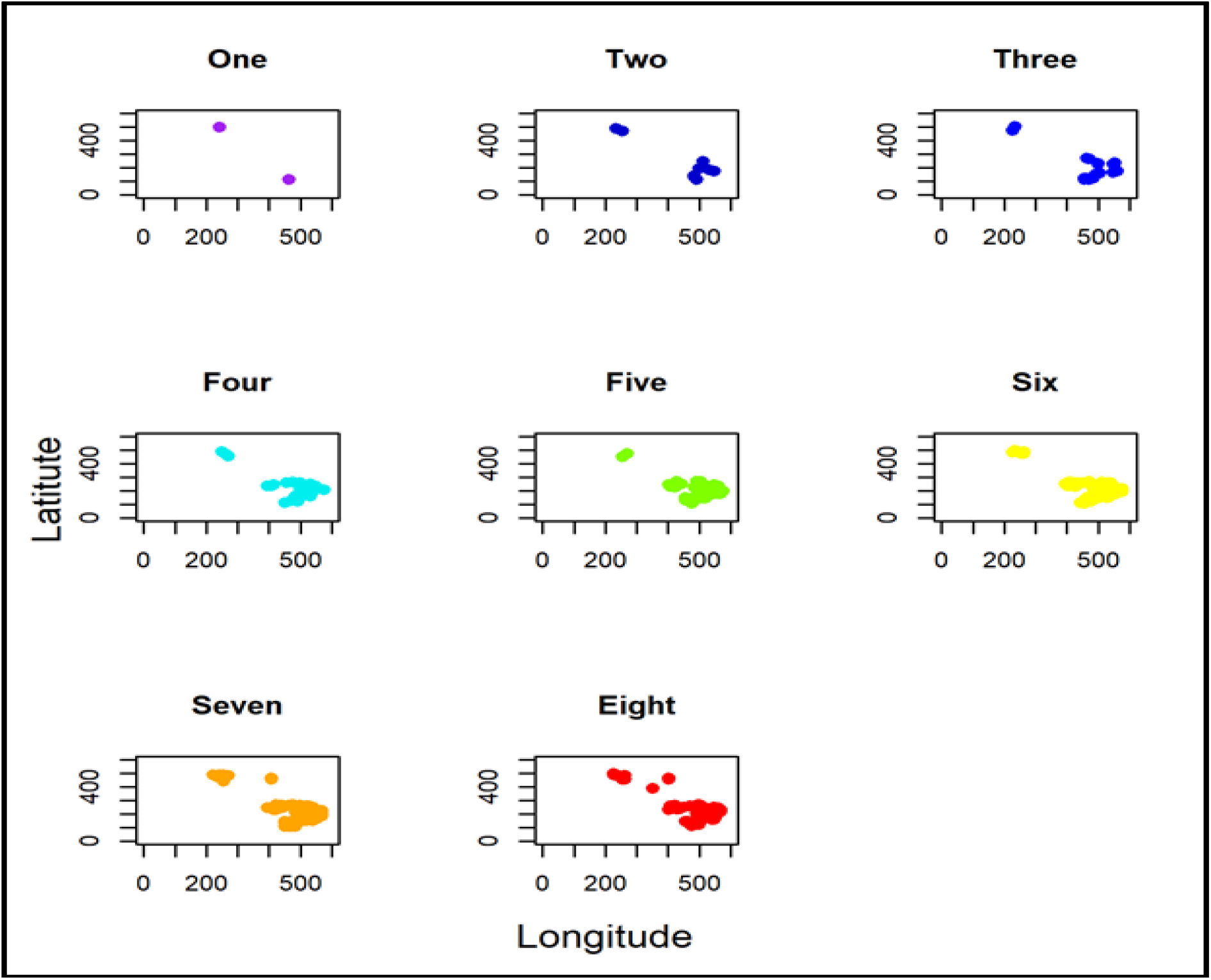
2D plots indicating the number of significant gene-sets across the coordination for COL11A1 at high CAF level

Finally, Figure 5 shows the combined significance *(CS)* heatmap of the 8 hallmarks for COL11A1 phenotype at high CAF level. Higher values of *CS* in Figure 5 represent locations with a combined significant gene-set.

**Figure 5.**
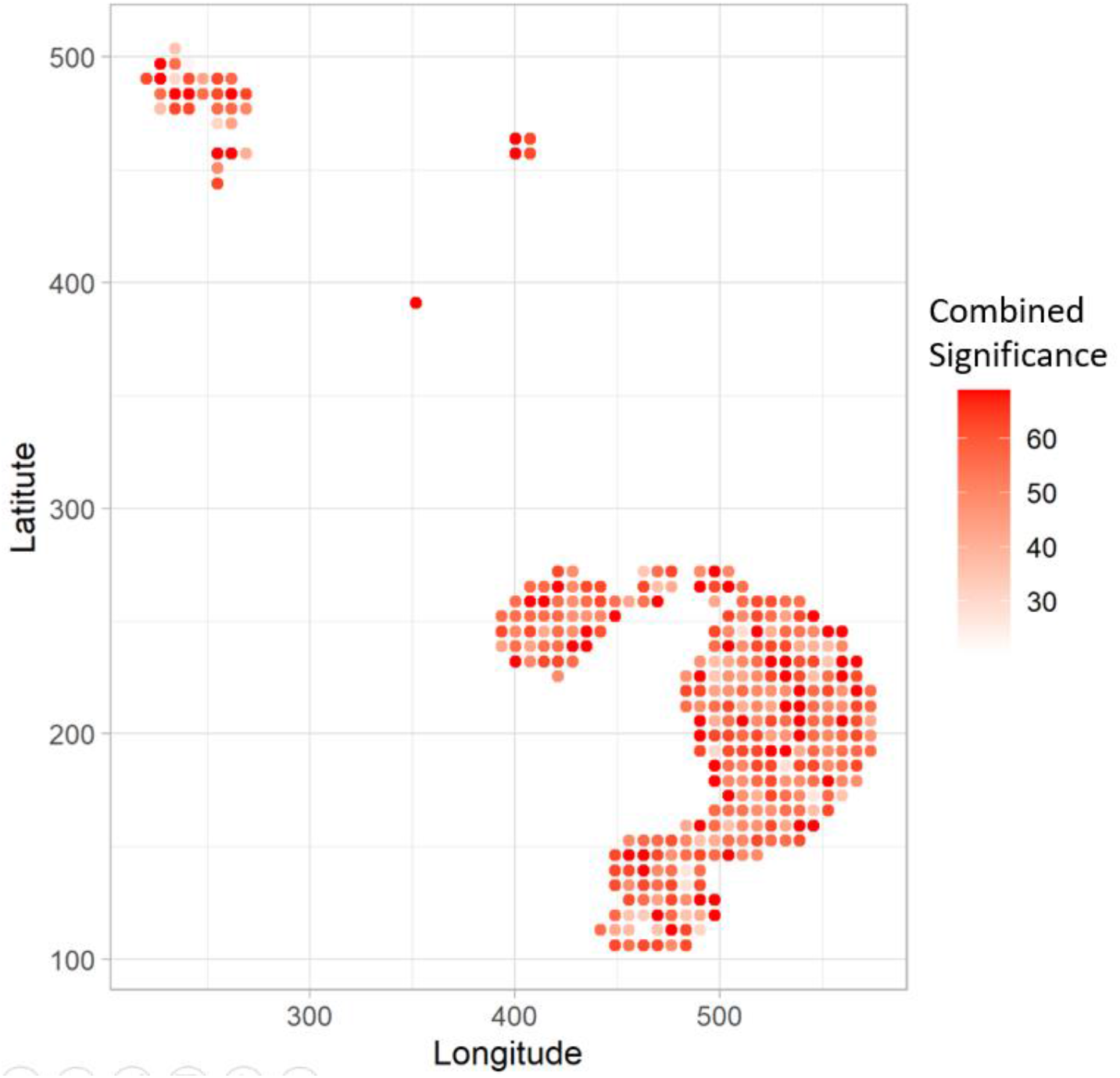
The spatial heatmap shows the Combined Significance (CS) of association of the 8 hallmarks with the COL11A1 phenotype at high CAF level. Bigger the value of CS at a location, the corresponding point is shown in darker red.

### Simulation study results

Table S1 and Figure 6 shows the estimated statistical power of the methods using the simulation study. GSEA has the least statistical power among the four methods at all the 144 scenarios. Using 500 iterations in the simulation study, it was revealed that regardless of any parameter in the simulation, GSEA and GWLCT always have the least and most statistical power among the methods, respectively. We note that bandwidth, number of coordinate points, gene-set size, or total number of genes in each simulation experiment, do not affect the statistical power performance of the methods, a desirable feature shared by sound gene-set analysis methods.

**Figure 6.**
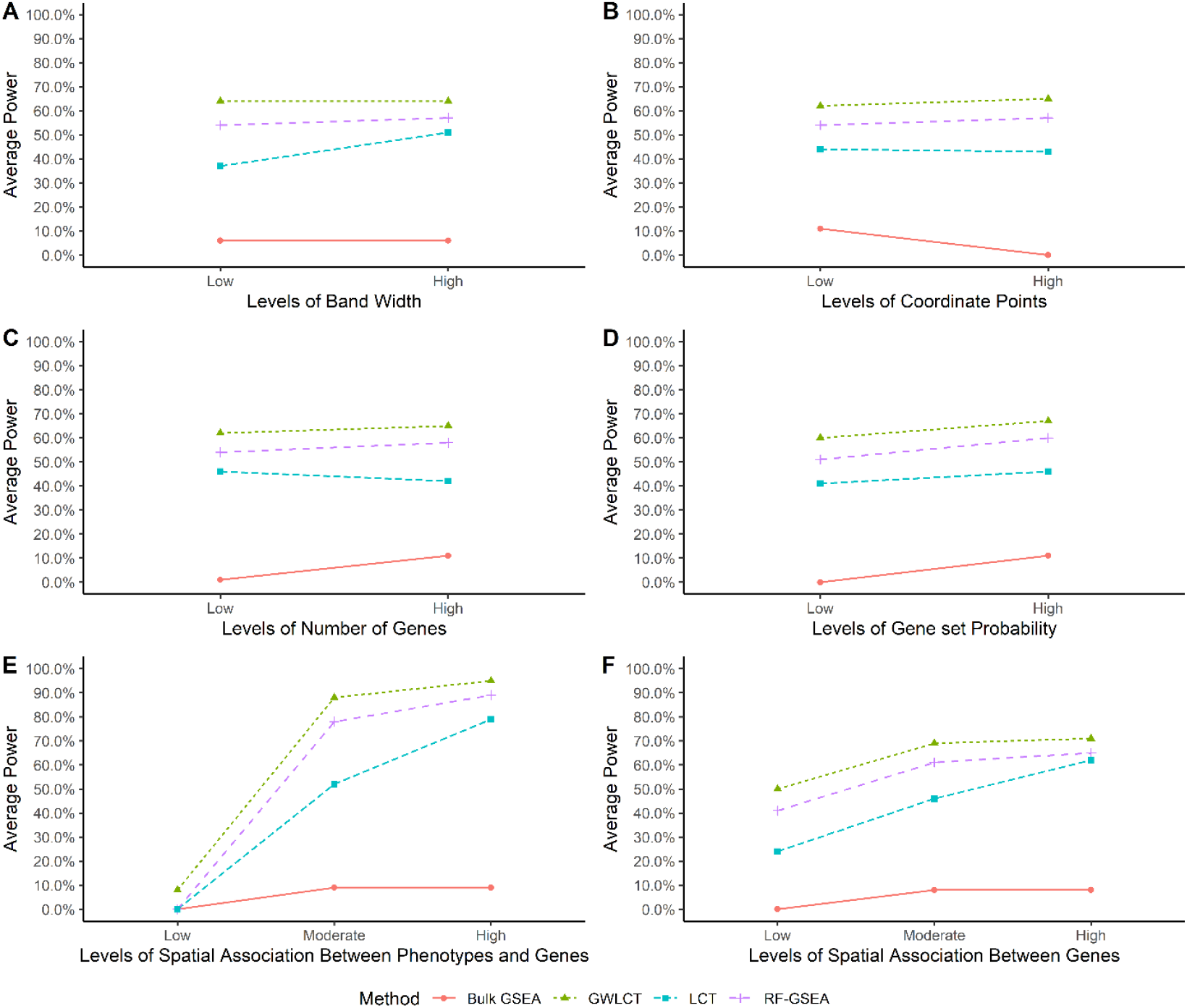
Comparing the average statistical power of each method across various levels of simulation parameters.

The GSEA statistical power is affected by all the simulation variables, however this is mostly due to the zero power of some scenarios. The performance of GSEA is affected by larger coordination, higher number of genes, higher spatial association among gene expressions, as well as between the phenotype and the gene expressions, and higher number of genes at the gene-sets leads to higher statistical power. Obviously, GSEA is in a lower class of statistical power compared to the other three approaches. Regardless of the scenarios, the results of one-way analysis of variance shows that there are significant differences in the statistical power of LCT, GWLCT, and RF-GSEA (F=7.842, p<0.001). Tukey’s multiple comparison revealed that the difference is due to lower statistical power of LCT compared to the other two methods. Considering a fixed effect for other variables in the simulation, one can find out that LCT has a lower statistical power compared to the other two GWLCT and RF-GSEA when the band width is low. For a study with a low number of coordination points, the power of GWLCT is significantly higher in comparison to LCT. The almost flat trend of power for the methods also reveals that the performance of the methods is robust against this parameter. As the number of genes increases, the power of LCT reduces significantly compared to GWLCT and RF-GSEA. Moreover, as the amount of spatial association among genes decreases, the power of LCT is significantly lower compared to GWLCT and RF-GSEA. At high levels of genes spatial association, LCT performs reasonably well compared to GWLCT and RF-GSEA, as it is designed to accommodate correlations across genes in a set or biological pathway, via a shrinkage correlation matrix. However, at lower levels of gene spatial correlation, GWLCT and RF-GSEA outperform LCT. Moreover, as the amount of spatial association between phenotype and genes increases, the power of GWLCT is significantly higher compared to LCT and RF-GSEA. In addition, the statistical power is robust against the probability of genes belonging to the gene-sets, which is directly related to gene-set size. Although the power increases slightly for all the methods when the size of gene-sets are larger, the increase is not statistically significant between Low and High levels of binomial probability parameter.

Therefore, the most important variables influencing statistical power are the spatial association features. There is a dose response trend of improvement in statistical power for each method, as spatial association among genes increases. The GWLCT outperforms all other methods in all levels of low, medium, and high spatial association among genes. The performance gap narrows down as we move from Low to High levels, indicating higher magnitudes of correlations are easier to be picked up. Interestingly, GWLCT picks up even on subtle spatial associations across genes, exhibiting the largest improvement in statistical power over other methods at Low spatial association levels. The spatial association between phenotype and gene expressions also plays a key role in method performance. When there is a low spatial association between phenotype and genes, GWLCT is still able to detect true significant associations between gene-sets and phenotype, while all other methods have a flat zero statistical power. Similar to the spatial associations across genes, GWLCT picks up on subtle signals for low levels of phenotype-gene expression levels of spatial associations. GWLCT outperforms all other methods at Moderate and High spatial association between phenotype and genes, with the highest statistical power of 0.95, across all simulation scenarios. A significant increase happens when the spatial association between phenotype and genes increases. The highest power for the GWLCT and RF-GSEA can be achieved when both spatial association variables are high. In contrast to GWLCT, the RF-GSEA loses its statistical power when the spatial association among the genes is at Low levels. GWLCT can identify more subtle signals of spatial association, which is an attractive property of the proposed method. More details on the statistical power of the methods in different circumstances can be found in the appendix Table S2.

## Discussion

Observations of heterogeneity of cell subpopulations in a tumor and the complex interplay of functions involved in the diverse morphological and phenotypic profiles of cancers have a long history. Even in the 19^th^ century, pleomorphism of cancer cells within tumors was observed by the “father of modern pathology”, Rudolph Virchow. More recently, in the 1970s, G.H. Heppner, I.J. Fidler and others showed the existence of distinct subpopulations of cancer cells in tumors, which differed in terms of their tumorigenicity, resistance to treatment, and potential to metastasize. Heppner reviewed the concept of tumor heterogeneity in 1984, and recognized cancers as being composed of multiple subpopulations (96), which leads to heterogeneity of cellular morphology, gene expression, metabolism, motility, proliferation, etc.(1). Importantly, ITH has been shown to be associated with poor outcome and decreased response to cancer treatment multiple human cancer types implying a general role in therapeutic resistance (5–7).

Gene-set analysis (GSA) is a well-established methodological approach in bioinformatics to test for significant regulation of a selected collection of genes across given samples that represent distinct outcomes. At the level of single cells, GSA could be extended to samples that are individual cells which admit to different phenotypes of interest (74). Furthermore, for spatial single cell analysis, such phenotypes would ideally have a spatially correlated and continuous representation. The gene-sets used in GSA are typically curated based on existing experimentally obtained knowledge of genes and their involvements in molecular pathways. In the present study, for illustrative purposes, we selected a collection of 8 gene-sets that represent certain distinctive hallmarks of cancer (97). To test for their enrichment in relevant intratumor contexts, we selected 5 different CAF phenotypes of interest since CAFs are well-known for their contribution to heterogeneity and plasticity in the tumor microenvironment (98).

The usual methods for GSA involve one of the two major approaches: (a) competitive, which examines if the correlation of a gene-set with the phenotype is the same as the other gene-sets, and (b) self-contained hypothesis, which investigates if the expression of a gene-set changes by the experimental condition. Our LCT method belongs to the former approach which is more likely than traditional methods to detect the regulation of a functional process or biological pathway that is significantly associated with the gene expression results of a given SCA experiment (74). Interestingly, LCT also extends to longitudinal (99), multivariate and continuous outcomes (72), which are capabilities that we built upon here for providing more accurate representation of single cell level stochasticity of the transcriptomic behavior than that of the univariate and discrete class labels typically used in traditional bulk sample studies.

Simulations, along with real omic data analysis, have served as a powerful and effective tool for establishing the performance of new GSA methods. Past studies have thus used simulation for comparative analysis of different criteria of performance of LCT and other major GSA methods. It was found that LCT has type I error and power that are comparable to MANOVA-GSA (72) and superior to SAM-GS (69), particularly at higher magnitudes of correlation values across gene-sets (as is commonly noted during GSA). In terms of computational efficiency, LCT outperformed both methods. In another simulation study, LCT also outperformed GSEA (100). Along this direction, therefore, in the present study, we conducted a large number of simulations to compare the performance of GWLCT against multiple known GSA techniques based on a variety of well-defined criteria under different experimental assumptions (or scenarios). Interestingly, statistical power did not change with a variation in set size, number of coordination points or bandwidth, or total number of genes in the simulation dataset, for any of the methods considered, which represent desirable properties for sound GSA methods. Larger spatial correlation across gene expression measurements and between genes and phenotype are key aspects of improved statistical power across our simulation experiments.

In the present study, we introduced GWLCT as a new computational platform that presents a fusion of ideas from spatial data analysis (GWR) and bioinformatics (GSA). We understand that the dual modes – both spatial and single-cell – in which GWLCT provides a joint extension to other GSA approaches places it in a unique category thus making it difficult to compare with the existing methods. Yet, we conducted extensive simulation studies which revealed better performance of GWLCT based on several criteria as compared to many known GSA methods that work either on bulk transcriptomics for different scenarios or aspatial version of single-cell transcriptomics. In particular, the use of multiple different kernels and flexible (adaptive) choice of corresponding bandwidths for geographical weighting allows the linear combination test to test for local associations between selected gene-sets and phenotypic contexts within a tumor sample. Thus, GWLCT provides a novel spatial version of gene-set analysis using high-resolution spatial scRNA-seq data. It does have some limitations that will be addressed in our future work. For instance, as it is difficult to determine a priori the precise spatial scale at which a gene-set or pathway may be regulated in a given phenotypic context, new multi-scale geographical weighting techniques (101) may prove to be useful. We will also extend GWLCT to other omic data as we have previously demonstrated with LCT (99). As spatial single cell omic platforms become increasingly popular, GWLCT will enrich the ongoing efforts in this rapidly emerging area of research (7, 55, 102).

## Acknowledgments

This research was supported by a Fellowship from Mathematics of Information Technology and Complex Systems Accelerate program.

## Contribution to the field

This study introduces a new computational platform, GWLCT, for combining gene set analysis with spatial data modeling to provide a new statistical test for association of continuous phenotypes and molecular pathways in spatial single-cell RNA-seq data collected from an input tumor sample. While the popular gene-set analysis (GSA) methods are aspatial and use only bulk gene expression data, GWLCT is developed for modeling with single cell gene expressions measured at given locations over a tumor space. The geographical weighting of GWLCT allows nearby neighborhoods to contribute more to each local model, and the regions with significant association of a selected gene-set and a corresponding phenotype are detected using scan statistics and illustrated as maps. The platform produces spatial heatmaps of combined significance over all selected gene-sets and also provides new 3D interactive tools for visualizing the associations in tumor space. In addition to illustrative analysis with real tumor data, extensive simulation studies demonstrate the superior performance of GWLCT over other methods. As spatial single cell omic platforms become increasingly popular, GWLCT will enrich the ongoing efforts in this rapidly emerging area of research.

## Availability of data and materials

The datasets used and/or analyzed during the current study would be available from the corresponding author on reasonable request.

## Author contributions

PA: Software coding, Data analysis, drafting the manuscript, reviewing the results and approving the final version of the manuscript

ID: study conception and design, interpreting the results, reviewing the results and approving the final version of the manuscript

SP: study conception and design, interpreting the results, reviewing the results and approving the final version of the manuscript

MH: Prepared the data, contributed data analysis, reviewing the results and approving the final version of the manuscript

## Ethics approval

Not applicable to our study as it uses publicly available anonymized data for secondary analysis.

