## Supplemental Tables for "Geographically Weighted Linear Combination Test for Gene Set Analysis of a Continuous Spatial Phenotype as applied to Intratumor Heterogeneity": Supplementary.pdf

### Supplementary Material:

**Supplementary table S1.** Mean (standard error) of statistical power for each method across levels of simulation parameters.

| **Simulation variables** | **Level** | **RF-GSEA** | **GSEA** | **Global LCT** | **GWLCT** |
|---|---|---|---|---|---|
| Kernel bandwidth | Low | 0.54(0.05) | 0.06(0.02) | 0.37(0.05) | 0.64(0.05) |
| | High | 0.57(0.05) | 0.06(0.03) | 0.51(0.05) | 0.64(0.05) |
| | P-value | 0.691 | 0.945 | 0.057 | 0.959 |
| Number of coordinate points | Low | 0.54(0.05) | 0.11(0.04) | 0.44(0.05) | 0.62(0.05) |
| | High | 0.57(0.05) | 0.00(0.00) | 0.43(0.05) | 0.65(0.05) |
| | P-value | 0.656 | 0.002 | 0.884 | 0.703 |
| Number of genes | Low | 0.54(0.05) | 0.01(0) | 0.46(0.05) | 0.62(0.05) |
| | High | 0.58(0.05) | 0.11(0.04) | 0.42(0.05) | 0.65(0.05) |
| | P-value | 0.586 | 0.007 | 0.598 | 0.639 |
| Gene spatial association | Low | **0.41(0.05)** | 0.00(0.00) | **0.24(0.05)** | **0.50(0.06)** |
| | Moderate | **0.61(0.06)** | 0.08(0.04) | **0.46(0.06)** | **0.69(0.06)** |
| | High | **0.65(0.07)** | 0.08(0.04) | **0.62(0.07)** | **0.71(0.06)** |
| | P-value | **0.013** | 0.122 | **0.001** | **0.028** |
| Phenotype-Gene spatial association | Low | **0.00(0.00)** | 0.00(0.00) | **0.00(0.00)** | **0.08(0.01)** |
| | Moderate | **0.78(0.04)** | 0.09(0.04) | **0.52(0.05)** | **0.88(0.03)** |
| | High | **0.89(0.03)** | 0.09(0.04) | **0.79(0.05)** | **0.95(0.02)** |
| | P-value | **<0.001** | 0.089 | **<0.001** | **<0.001** |
| Gene-set probability | Low | 0.51(0.05) | 0.00(0.00) | 0.41(0.05) | 0.60(0.05) |
| | High | 0.60(0.05) | 0.11(0.04) | 0.46(0.05) | 0.67(0.05) |
| | P-value | 0.231 | 0.002 | 0.489 | 0.293 |

| **Supplementary Table S2.** Statistical power of the performed methods using 144 scenarios in the simulation study | | | | | | | | | | |
|---|---|---|---|---|---|---|---|---|---|---|
| **Scenario** | **Bandwidth** | **Coordination points** | **No Genes** | **Gene SA** | **Phenotype Gene SA** | **Gene set probability** | **RF-GSEA** | **GSEA** | **LCT** | **GWLCT** |
| 1 | Low | Low | Low | High | High | Low | 1 | 0 | 1 | 1 |
| 2 | High | Low | Low | High | High | Low | 1 | 0 | 1 | 1 |
| 3 | Low | Low | High | High | High | Low | 1 | 0 | 1 | 1 |
| 4 | High | Low | High | High | High | Low | 1 | 0 | 1 | 1 |
| 5 | Low | Low | Low | High | High | High | 1 | 0.0909 | 1 | 1 |
| 6 | High | Low | Low | High | High | High | 1 | 0 | 1 | 1 |
| 7 | Low | Low | High | High | High | High | 1 | 0.9091 | 1 | 1 |
| 8 | High | Low | High | High | High | High | 1 | 1 | 1 | 1 |
| 9 | Low | Low | Low | Moderate | High | Low | 1 | 0 | 0.792 | 1 |
| 10 | High | Low | Low | Moderate | High | Low | 1 | 0 | 1 | 1 |
| 11 | Low | Low | High | Moderate | High | Low | 0.9636 | 0 | 0.8371 | 1 |
| 12 | High | Low | High | Moderate | High | Low | 1 | 0 | 1 | 1 |
| 13 | Low | Low | Low | Moderate | High | High | 1 | 0.0909 | 0.8545 | 1 |
| 14 | High | Low | Low | Moderate | High | High | 1 | 0 | 1 | 1 |
| 15 | Low | Low | High | Moderate | High | High | 1 | 0.9091 | 0.8611 | 1 |
| 16 | High | Low | High | Moderate | High | High | 1 | 1 | 1 | 1 |
| 17 | Low | Low | Low | Low | High | Low | 0.2333 | 0 | 0.252 | 0.432 |
| 18 | High | Low | Low | Low | High | Low | 0.2 | 0 | 0.2 | 0.365 |
| 19 | Low | Low | High | Low | High | Low | 0.5455 | 0 | 0.0073 | 0.745 |
| 20 | High | Low | High | Low | High | Low | 0.6 | 0 | 0.6 | 0.875 |
| 21 | Low | Low | Low | Low | High | High | 1 | 0.0909 | 0.4102 | 1 |
| 22 | High | Low | Low | Low | High | High | 1 | 0 | 1 | 1 |
| 23 | Low | Low | High | Low | High | High | 1 | 0 | 0.0065 | 1 |
| 24 | High | Low | High | Low | High | High | 1 | 0 | 1 | 1 |

| | | | | | | | | | | |
|---|---|---|---|---|---|---|---|---|---|---|
| 25 | Low | High | Low | High | High | Low | 1 | 0 | 1 | 1 |
| 26 | High | High | Low | High | High | Low | 1 | 0 | 1 | 1 |
| 27 | Low | High | High | High | High | Low | 1 | 0 | 1 | 1 |
| 28 | High | High | High | High | High | Low | 1 | 0 | 1 | 1 |
| 29 | Low | High | Low | High | High | High | 1 | 0 | 1 | 1 |
| 30 | High | High | Low | High | High | High | 1 | 0 | 1 | 1 |
| 31 | Low | High | High | High | High | High | 1 | 0 | 1 | 1 |
| 32 | High | High | High | High | High | High | 1 | 0 | 1 | 1 |
| 33 | Low | High | Low | Moderate | High | Low | 1 | 0 | 0.8322 | 1 |
| 34 | High | High | Low | Moderate | High | Low | 1 | 0 | 1 | 1 |
| 35 | Low | High | High | Moderate | High | Low | 1 | 0 | 0.878 | 1 |
| 36 | High | High | High | Moderate | High | Low | 1 | 0 | 1 | 1 |
| 37 | Low | High | Low | Moderate | High | High | 1 | 0 | 0.9111 | 1 |
| 38 | High | High | Low | Moderate | High | High | 1 | 0 | 1 | 1 |
| 39 | Low | High | High | Moderate | High | High | 1 | 0 | 0.9653 | 1 |
| 40 | High | High | High | Moderate | High | High | 1 | 0 | 1 | 1 |
| 41 | Low | High | Low | Low | High | Low | 0.4222 | 0 | 0.1467 | 0.6 |
| 42 | High | High | Low | Low | High | Low | 0.4 | 0 | 0.216 | 0.6 |
| 43 | Low | High | High | Low | High | Low | 0.7455 | 0 | 0.1782 | 0.9636 |
| 44 | High | High | High | Low | High | Low | 0.6 | 0 | 0.4 | 1 |
| 45 | Low | High | Low | Low | High | High | 1 | 0 | 0.618 | 1 |
| 46 | High | High | Low | Low | High | High | 1 | 0 | 1 | 1 |
| 47 | Low | High | High | Low | High | High | 1 | 0 | 0.18 | 1 |
| 48 | High | High | High | Low | High | High | 1 | 0 | 0.8 | 1 |
| 49 | Low | Low | Low | High | Moderate | Low | 0.8333 | 0 | 0.8733 | 1 |
| 50 | High | Low | Low | High | Moderate | Low | 1 | 0 | 1 | 1 |
| 51 | Low | Low | High | High | Moderate | Low | 0.9091 | 0 | 0.3273 | 1 |
| 52 | High | Low | High | High | Moderate | Low | 1 | 0 | 1 | 1 |
| 53 | Low | Low | Low | High | Moderate | High | 0.9091 | 0.0909 | 0.7491 | 1 |
| 54 | High | Low | Low | High | Moderate | High | 1 | 0 | 1 | 1 |

| | | | | | | | | | | |
|---|---|---|---|---|---|---|---|---|---|---|
| 55 | Low | Low | High | High | Moderate | High | 0.9091 | 0.9091 | 0.3273 | 0.9091 |
| 56 | High | Low | High | High | Moderate | High | 1 | 1 | 1 | 1 |
| 57 | Low | Low | Low | Moderate | Moderate | Low | 0.8333 | 0 | 0.7027 | 0.9 |
| 58 | High | Low | Low | Moderate | Moderate | Low | 1 | 0 | 1 | 1 |
| 59 | Low | Low | High | Moderate | Moderate | Low | 0.9091 | 0 | 0.0218 | 0.9818 |
| 60 | High | Low | High | Moderate | Moderate | Low | 1 | 0 | 0.992 | 1 |
| 61 | Low | Low | Low | Moderate | Moderate | High | 0.3636 | 0.0909 | 0.4822 | 1 |
| 62 | High | Low | Low | Moderate | Moderate | High | 0.8 | 0 | 0.6 | 1 |
| 63 | Low | Low | High | Moderate | Moderate | High | 0.9455 | 0.9091 | 0.0145 | 0.9636 |
| 64 | High | Low | High | Moderate | Moderate | High | 1 | 1 | 0.984 | 1 |
| 65 | Low | Low | Low | Low | Moderate | Low | 0.1667 | 0 | 0.1707 | 0.2 |
| 66 | High | Low | Low | Low | Moderate | Low | 0.2 | 0 | 0.2 | 0.3512 |
| 67 | Low | Low | High | Low | Moderate | Low | 0.3455 | 0 | 0 | 0.5636 |
| 68 | High | Low | High | Low | Moderate | Low | 0.4 | 0 | 0.416 | 0.5629 |
| 69 | Low | Low | Low | Low | Moderate | High | 0.5455 | 0.0909 | 0.2036 | 0.6566 |
| 70 | High | Low | Low | Low | Moderate | High | 0.6 | 0 | 0 | 0.724 |
| 71 | Low | Low | High | Low | Moderate | High | 0.6455 | 0 | 0 | 0.7818 |
| 72 | High | Low | High | Low | Moderate | High | 0.6 | 0 | 0.928 | 0.8 |
| 73 | Low | High | Low | High | Moderate | Low | 1 | 0 | 0.7467 | 1 |
| 74 | High | High | Low | High | Moderate | Low | 1 | 0 | 1 | 1 |
| 75 | Low | High | High | High | Moderate | Low | 1 | 0 | 0.9091 | 1 |
| 76 | High | High | High | High | Moderate | Low | 1 | 0 | 0.98 | 1 |
| 77 | Low | High | Low | High | Moderate | High | 1 | 0 | 0.7636 | 1 |
| 78 | High | High | Low | High | Moderate | High | 1 | 0 | 1 | 1 |
| 79 | Low | High | High | High | Moderate | High | 1 | 0 | 0.9091 | 1 |
| 80 | High | High | High | High | Moderate | High | 1 | 0 | 0.98 | 1 |
| 81 | Low | High | Low | Moderate | Moderate | Low | 0.9778 | 0 | 0.1631 | 1 |
| 82 | High | High | Low | Moderate | Moderate | Low | 0.8 | 0 | 0.512 | 1 |
| 83 | Low | High | High | Moderate | Moderate | Low | 0.8182 | 0 | 0.3665 | 1 |
| 84 | High | High | High | Moderate | Moderate | Low | 0.8 | 0 | 0.098 | 1 |

| | | | | | | | | | | |
|---|---|---|---|---|---|---|---|---|---|---|
| 85 | Low | High | Low | Moderate | Moderate | High | 1 | 0 | 0.027 | 1 |
| 86 | High | High | Low | Moderate | Moderate | High | 1 | 0 | 0.21 | 1 |
| 87 | Low | High | High | Moderate | Moderate | High | 1 | 0 | 0.5566 | 1 |
| 88 | High | High | High | Moderate | Moderate | High | 1 | 0 | 0.202 | 1 |
| 89 | Low | High | Low | Low | Moderate | Low | 0.208 | 0 | 0.4272 | 0.4 |
| 90 | High | High | Low | Low | Moderate | Low | 0.4 | 0 | 0.276 | 0.4 |
| 91 | Low | High | High | Low | Moderate | Low | 0.5778 | 0 | 0.0963 | 0.5778 |
| 92 | High | High | High | Low | Moderate | Low | 0.4 | 0 | 0 | 0.6 |
| 93 | Low | High | Low | Low | Moderate | High | 0.9926 | 0 | 0.7858 | 1 |
| 94 | High | High | Low | Low | Moderate | High | 1 | 0 | 0.698 | 1 |
| 95 | Low | High | High | Low | Moderate | High | 0.8074 | 0 | 0.0982 | 1 |
| 96 | High | High | High | Low | Moderate | High | 0.9256 | 0 | 0.1232 | 1 |
| 97 | Low | Low | Low | High | Low | Low | 0 | 0 | 0.002 | 0.235 |
| 98 | High | Low | Low | High | Low | Low | 0 | 0 | 0.0002 | 0.1565 |
| 99 | Low | Low | High | High | Low | Low | 0 | 0 | 0.0012 | 0.2155 |
| 100 | High | Low | High | High | Low | Low | 0 | 0 | 0.0021 | 0.1321 |
| 101 | Low | Low | Low | High | Low | High | 0 | 0 | 0.002 | 0.1985 |
| 102 | High | Low | Low | High | Low | High | 0 | 0 | 0.0015 | 0.1365 |
| 103 | Low | Low | High | High | Low | High | 0 | 0 | 0.009 | 0.0985 |
| 104 | High | Low | High | High | Low | High | 0 | 0 | 0.0022 | 0.0985 |
| 105 | Low | Low | Low | Moderate | Low | Low | 0 | 0 | 0.0032 | 0.118 |
| 106 | High | Low | Low | Moderate | Low | Low | 0 | 0 | 0.0042 | 0.1049 |
| 107 | Low | Low | High | Moderate | Low | Low | 0 | 0 | 0.0037 | 0.1024 |
| 108 | High | Low | High | Moderate | Low | Low | 0 | 0 | 0.0026 | 0.0989 |
| 109 | Low | Low | Low | Moderate | Low | High | 0 | 0 | 0 | 0.0894 |
| 110 | High | Low | Low | Moderate | Low | High | 0 | 0 | 0 | 0.0941 |
| 111 | Low | Low | High | Moderate | Low | High | 0 | 0 | 0.0001 | 0.0998 |
| 112 | High | Low | High | Moderate | Low | High | 0 | 0 | 0 | 0.1 |
| 113 | Low | Low | Low | Low | Low | Low | 0 | 0 | 0 | 0.007 |
| 114 | High | Low | Low | Low | Low | Low | 0 | 0 | 0 | 0.0006 |

| | | | | | | | | | | |
|---|---|---|---|---|---|---|---|---|---|---|
| 115 | Low | Low | High | Low | Low | Low | 0 | 0 | 0 | 0.0006 |
| 116 | High | Low | High | Low | Low | Low | 0 | 0 | 0.0002 | 0.0006 |
| 117 | Low | Low | Low | Low | Low | High | 0 | 0 | 0 | 0.0002 |
| 118 | High | Low | Low | Low | Low | High | 0 | 0 | 0 | 0.0001 |
| 119 | Low | Low | High | Low | Low | High | 0 | 0 | 0.0001 | 0.0004 |
| 120 | High | Low | High | Low | Low | High | 0 | 0 | 0 | 0.0004 |
| 121 | Low | High | Low | High | Low | Low | 0 | 0 | 0.0002 | 0.1416 |
| 122 | High | High | Low | High | Low | Low | 0 | 0 | 0.0092 | 0.1322 |
| 123 | Low | High | High | High | Low | Low | 0 | 0 | 0.0004 | 0.1654 |
| 124 | High | High | High | High | Low | Low | 0 | 0 | 0.0004 | 0.0917 |
| 125 | Low | High | Low | High | Low | High | 0 | 0 | 0.0009 | 0.1716 |
| 126 | High | High | Low | High | Low | High | 0 | 0 | 0.0015 | 0.1157 |
| 127 | Low | High | High | High | Low | High | 0 | 0 | 0.009 | 0.1027 |
| 128 | High | High | High | High | Low | High | 0 | 0 | 0.0022 | 0.0872 |
| 129 | Low | High | Low | Moderate | Low | Low | 0 | 0 | 0 | 0.0817 |
| 130 | High | High | Low | Moderate | Low | Low | 0 | 0 | 0 | 0.0991 |
| 131 | Low | High | High | Moderate | Low | Low | 0 | 0 | 0 | 0.0813 |
| 132 | High | High | High | Moderate | Low | Low | 0 | 0 | 0 | 0.102 |
| 133 | Low | High | Low | Moderate | Low | High | 0 | 0 | 0 | 0.0917 |
| 134 | High | High | Low | Moderate | Low | High | 0 | 0 | 0 | 0.0715 |
| 135 | Low | High | High | Moderate | Low | High | 0 | 0 | 0 | 0.0709 |
| 136 | High | High | High | Moderate | Low | High | 0 | 0 | 0 | 0.1085 |
| 137 | Low | High | Low | Low | Low | Low | 0 | 0 | 0 | 0 |
| 138 | High | High | Low | Low | Low | Low | 0 | 0 | 0 | 0 |
| 139 | Low | High | High | Low | Low | Low | 0 | 0 | 0 | 0 |
| 140 | High | High | High | Low | Low | Low | 0 | 0 | 0 | 0 |
| 141 | Low | High | Low | Low | Low | High | 0 | 0 | 0 | 0 |
| 142 | High | High | Low | Low | Low | High | 0 | 0 | 0 | 0 |
| 143 | Low | High | High | Low | Low | High | 0 | 0 | 0 | 0 |
| 144 | High | High | High | Low | Low | High | 0 | 0 | 0 | 0 |

SA: Spatial association; GSEA: Gene Set Enrichment Analysis; LCT: Linear combination test; GWLCT: geographically weighted linear combination test
